# Comprehensive skin microbiome analysis reveals the uniqueness of human-associated microbial communities among the class Mammalia

**DOI:** 10.1101/201434

**Authors:** Ashley A. Ross, Kirsten Müller, J. Scott Weese, Josh D. Neufeld

**Affiliations:** Department of Biology, University of Waterloo, Waterloo, Ontario, Canada; Department of Pathobiology, University of Guelph, Guelph, Ontario, Canada

## Abstract

Skin is the largest organ of the body and represents the primary physical barrier between mammals and their external environment. The objective of this research was to generate a skin microbiota baseline for members of the class Mammalia, testing the effects of host species, geographic location, body region, and biological sex. The back, torso, and inner thigh regions of 177 non-human mammals were collected to include representatives from 38 species and 10 mammalian orders. Animals were collected from local farms, zoos, households, and the wild. All samples were amplified using the V3-V4 16S rRNA gene region and sequenced using a MiSeq (Illumina). For reference, previously published skin microbiome data from 20 human participants, sampled using an identical protocol to the non-human mammals, were included in the analysis. Human skin was significantly less diverse than all other mammalian orders and the factor most strongly associated with community variation for all samples was whether the host was a human. Within non-human samples, host taxonomic order was the most significant factor influencing the skin community, followed by the geographic location of the habitat. By comparing the congruence between known host phylogeny and microbial community dendrograms, we observed that Artiodactyla (even-toed ungulates) and Perissodactyla (odd-toed ungulates) had significant congruence, providing first evidence of phylosymbiosis between skin communities and their hosts.

**Significance:** Skin forms a critical protective barrier between a mammal and its external environment. Baseline data on the mammalian skin microbiome is crucial for making informed decisions related to veterinary research and biodiversity conservation strategies, in addition to providing insight into mammalian evolutionary history. To our knowledge, this study represents the largest mammalian skin microbiota project to date. These findings demonstrate that human skin is distinct, not only from other Primates, but from all 10 mammalian orders sampled. Using phylosymbiosis analysis, we provide the first evidence that co-evolution may be occurring between skin communities and their mammalian hosts, which warrants more in-depth future studies of the relationships between mammals and their skin microbiota.

## Introduction

Skin is the largest organ of the body and represents the primary physical barrier between mammals and their external environment. Characterization of the microbiota on skin is essential for diagnosing skin conditions (1), understanding how an animal coevolves with its microbiota (2), and studying the interactions between microbiota and the host immune system (3). Skin microorganisms also produce compounds that influence animal behavior, such as intra-specific behavior modifying pheromones (4) and volatile organic compounds resulting in body odour (5–7). Cultivated human skin microbiota have been linked to the rates at which hosts are bitten by mosquitos (8, 9), which has implications for disease transmission, such as malaria.

High-throughput sequencing has provided the ability to evaluate factors that influence the skin microbiota and how these microbial communities impact health and skin conditions. Humans are uniquely colonized by skin microbial communities that vary between body regions (10–12), individuals (13), age (14, 15), and diet (16). Skin conditions, such as atopic dermatitis, can occur when the resident skin microbial community undergoes dysbiosis, which is defined as a community shift from the normal microbiota (17, 18). The composition of human skin microbial communities can also be linked to host hygiene. Previous studies have shown that skin microbial communities are affected by deodorants, soaps, and cosmetics (19–21). Indeed, three dimensional maps of human skin have shown that the residues from skin products are detectable and can influence the skin microbiota (12).

Although many studies have characterized the human microbiome, far less is known about the skin microbiome of non-human mammals, especially from studies that employed high-throughput sequencing techniques. Previous skin microbiome studies of cats (22–24) and dogs (25) demonstrated that cats with allergies have higher levels of the fungi *Agaricomycetes* and *Sordariomycetes* (22) and bacterial communities that are unique to individual felines (24), whereas allergic dogs exhibited lower bacterial species richness. Moreover, bovine skin afflicted with bovine digital dermatitis possesses a distinct microbiota from healthy skin (26). Hence, a baseline dataset of what constitutes healthy skin microbiota for a variety of species is crucial for determining the cause of skin ailments.

Multiple studies have been conducted on both wild and captive animals to elucidate the roles of host species, geographic location, body region, and captivity status exhibit on the skin microbiota (24, 25, 27–30). Analyzing skin samples of 63 individuals from five primate species revealed that human axillae were associated with distinct microbial communities, with lower overall diversity (27). The authors suggested that differences were due both to human hygiene, as well as host-microbe evolution. Skin biopsies and sloughed skin from 56 free-swimming humpback whales (*Megaptera novaeangliae*) from the North Pacific, South Pacific, and North Atlantic oceans demonstrated that core genera were present despite large geographical distances (29). However, skin microbial communities also exhibited shifts between geographic locations and whale satiation states throughout their migration. A large study of bats determined that the host species, geographic region, and site were significant factors influencing skin communities (28). Microbial diversity of skin and pouch samples from Tasmanian devils demonstrated a strong influence of geographic location and revealed significant differences between wild and captive populations (31). There was a significant difference in the abundance of 159 skin OTUs between the wild and captive groups, such as absence of *Brochothrix* and increase in *Mycobacterium* in captive Tasmanian devils. Together, these previous studies indicate that both phylogeny and habitat can impact skin microbial communities. However, studies that focus on a wide range of species and body location are important for better elucidating factors that influence skin microbial communities and generating a clear understanding of microbiota-host coevolution, especially considering that species sampled to date represent only a small proportion of known mammals.

The objective of this research was to generate a preliminary skin microbiome dataset for the class Mammalia in Southern Ontario, Canada in order to identify correlations of skin microbiota with species, geographical location, hygiene, and body region. Comprehensive understanding of the mammalian skin microbiome is crucial for making informed decisions related to veterinarian research and conservation strategies, with implications for mammalian evolutionary history.

## Results and Discussion

The diversity of bacteria and archaea were analyzed from 589 mammalian skin samples (**Table 1**). A total of 22,728 unique operational taxonomic units (OTUs) were obtained that corresponded to 44 prokaryotic phyla. The following general taxonomic distributions of the mammalian skin microbiota exclude the human samples that were published previously as part of a broader human cohabitation study (32). There were six phyla present above 1% abundance that constituted a mean of 96.0±4.0% of all reads (Supplementary Table 1): *Firmicutes* (33.6±20.4%), *Proteobacteria* (28.5±19.1%), *Actinobacteria* (23.6±16.1%), *Bacteroidetes* (7.6±4.9%), *Cyanobacteria* (1.5±2.6%), and *Chloroflexi* (1.1±1.8%). These abundances represent a significant decrease of 33.2% in the abundance of the phyla *Actinobacteria* (*p* < 0.001), and significant increases in the abundance of *Bacteroidetes* (*p* < 0.001), *Chloroflexi* (*p* < 0. 001), *Cyanobacteria* (*p* = 0.01), and *Proteobacteria* (*p* < 0.001) compared to human skin samples of the same body regions. The OTU with the highest relative abundance varied among species (Table 1).

**Table 1:**
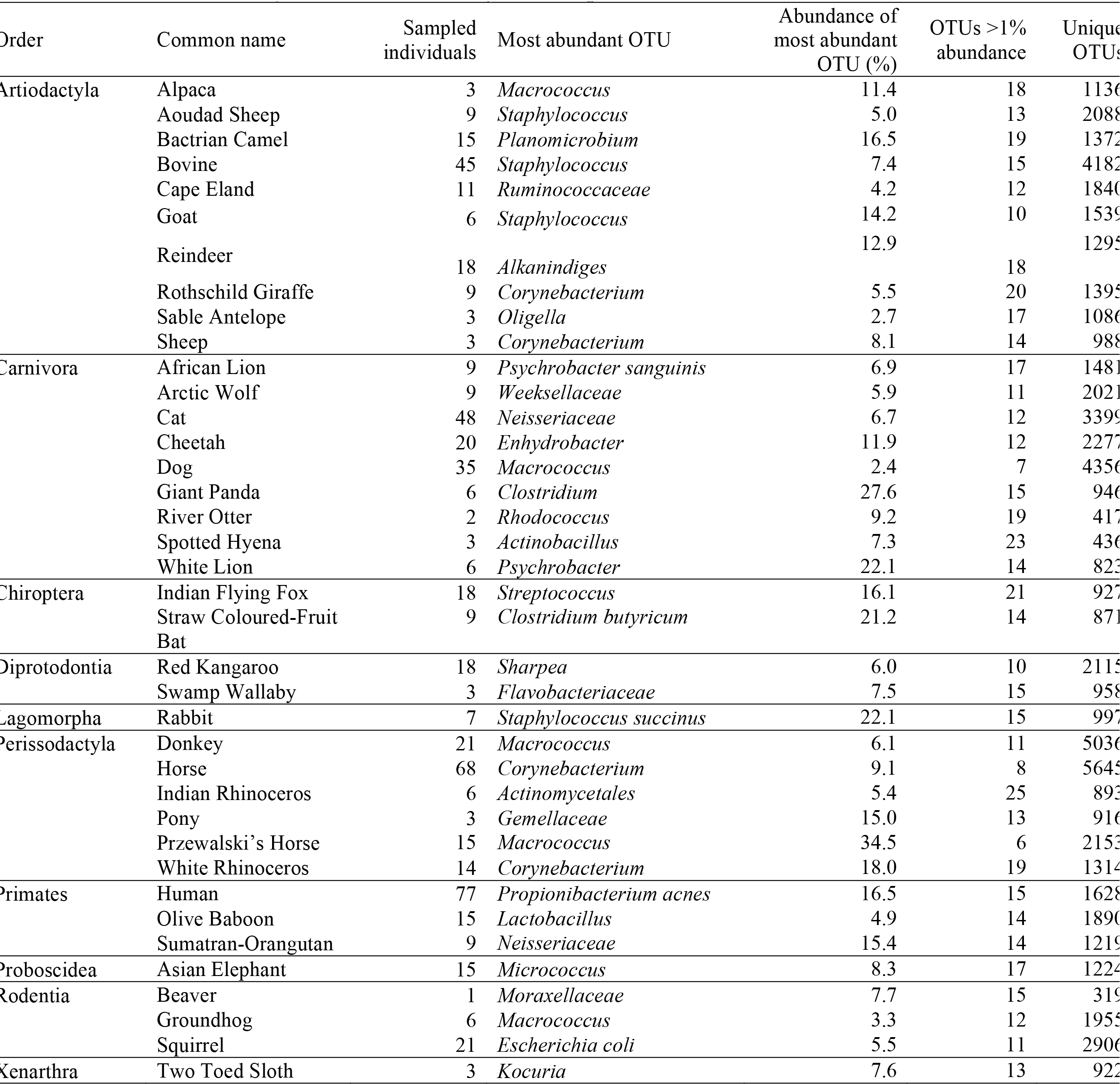
Summary of each mammalian species sampled with data on the most abundant OTU.

### Humans have a distinct microbial community from the majority of animals

An indicator species analysis determined that all human samples had elevated levels of *S. epidermidis*, *Corynebacterium*, and *Propionibacterium acnes* (Table 2), which is in accordance with previous literature (10, 33, 34). In contrast, non-human animals (“animals”) were associated with soil-related organisms, such as *Arthrobacter* and *Sphingomonas*, albeit at lower average abundance than human indicators. This finding was corroborated by a core OTU analysis ( Supplementary Figure 1). A core OTU was defined as one that was present in a minimum of 90% of non-rarefied samples. All mammalian clades shared six core OTUs including *Arthrobacter*, *Sphingomonas*, and *Agrobacterium.* Five mammalian orders were analyzed further that contained multiple host species and were not composed of cats, dogs, or humans. Each of the orders except Perissodactyla had core OTUs that were not shared with any of the other mammalian orders. These core OTUs represent microbiota that persist despite different varying geographical locations and enclosures. The presence of a large proportion of soil organisms (Supplementary Figure 2) may be explained by frequent contact of the skin with the external environment. Although the sampling of terrestrial mammals did not include a step to rinse off environmental microbiota, as has been completed in amphibian microbiota studies (35), future studies might test alternative sampling protocols to access the mammalian skin microbiota in order to minimize sampling of allochthonous microorganisms. Resident and transient microbiota on skin has been discussed previously for human clinical applications (36, 37). Medically, the organism is transient if it can easily be disinfected with antiseptics, or removed with soap and water. Environmental influences, such as the observed soil organisms, are expected components of the transient microbiota. The sampling methods used in this study would detect both the resident and transient microbiota.

**Table 2:**
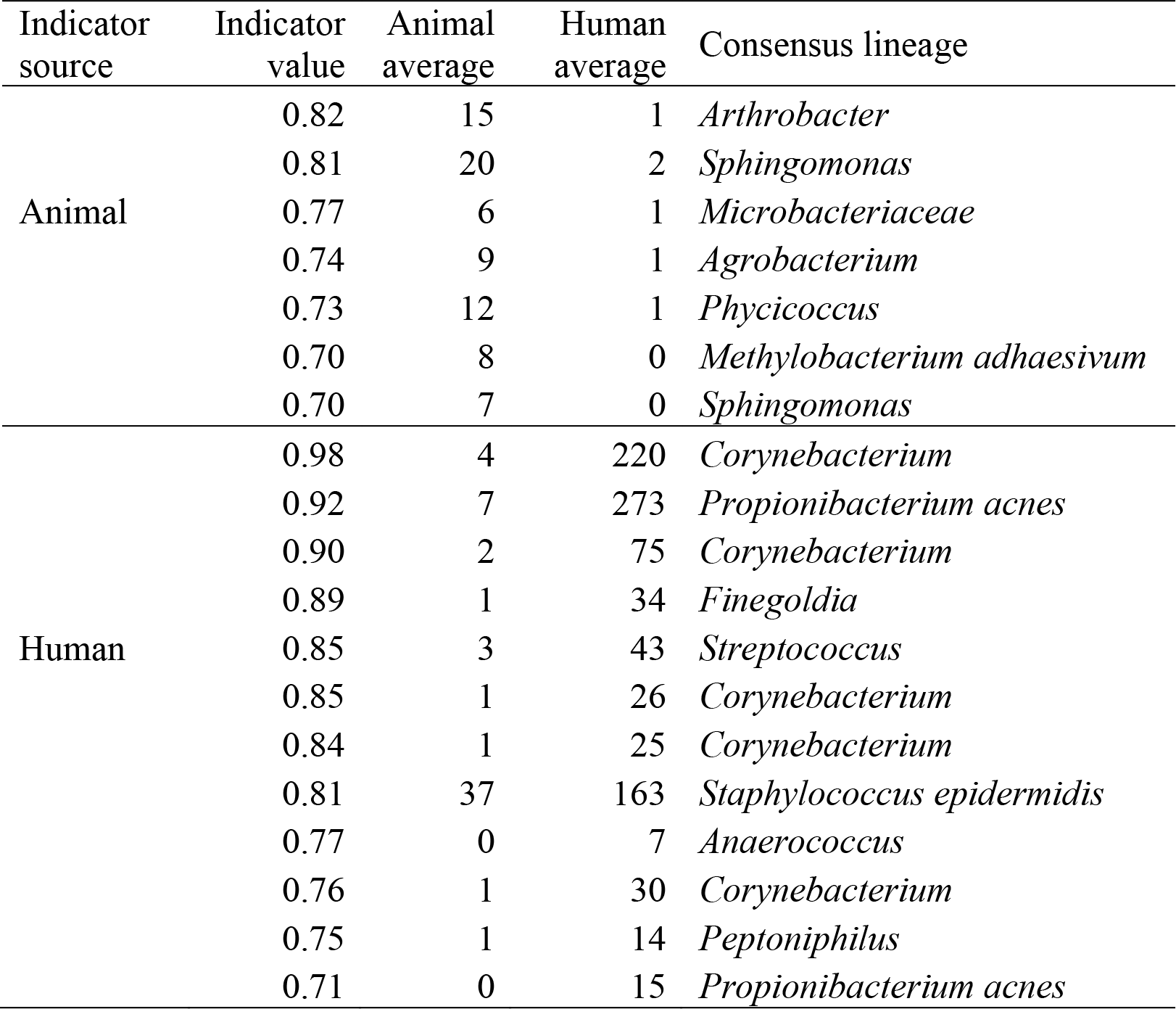
Indicator analysis of human and non-human animals. Indicator OTUs were defined as having an indicator value threshold of >0.7 and *p* < 0.05. Reported averages correspond to the number of sequences per sample, rarefied to 1654 reads per sample total. Multiple OTUs with the same genus are different strains.

Human samples possessed a unique microbial community from all other non-human mammals, except for several domestic pets from the order Carnivora (Figure 1). In addition, human skin was significantly less diverse than all other mammalian orders, according to both the number of distinct OTUs (Figure 2A) and Shannon indices (6.54 vs 3.96, *p* < 0.001; Figure 2B), which supports the findings of a study on five primate species that determined humans have lower diversity than other primates (27). Other orders whose microbial communities grouped tightly together include Diprotodontia (kangaroos), Chiroptera (bats), Rodentia (squirrels), and non-human Primates. A subsequent PERMANOVA analysis demonstrated that the factor most strongly associated with community variation for all samples was whether the host was a human (*F_1,587_* = 37.8, *p* < 0.001; Figure 3). Because humans have undergone recent evolutionary divergence from other non-human primates, such as orangutans (12-15 MYA) and baboons (21-25 MYA) (38), these results suggest that modern human practices, such as spending the majority of time indoors, frequent bathing, and wearing clothing may have impacted the diversity and composition of measured skin microbiota. A portion of the higher diversity in non-human mammals may be due to an increase in the number of transient microbiota. However, a study of previously uncontacted Amerindians demonstrated that changes in lifestyle, such as living outdoors, resulted in higher diversity (39), lending further support to this finding that modern human practices may be rapidly changing the skin microbiota.

**Figure 1:**
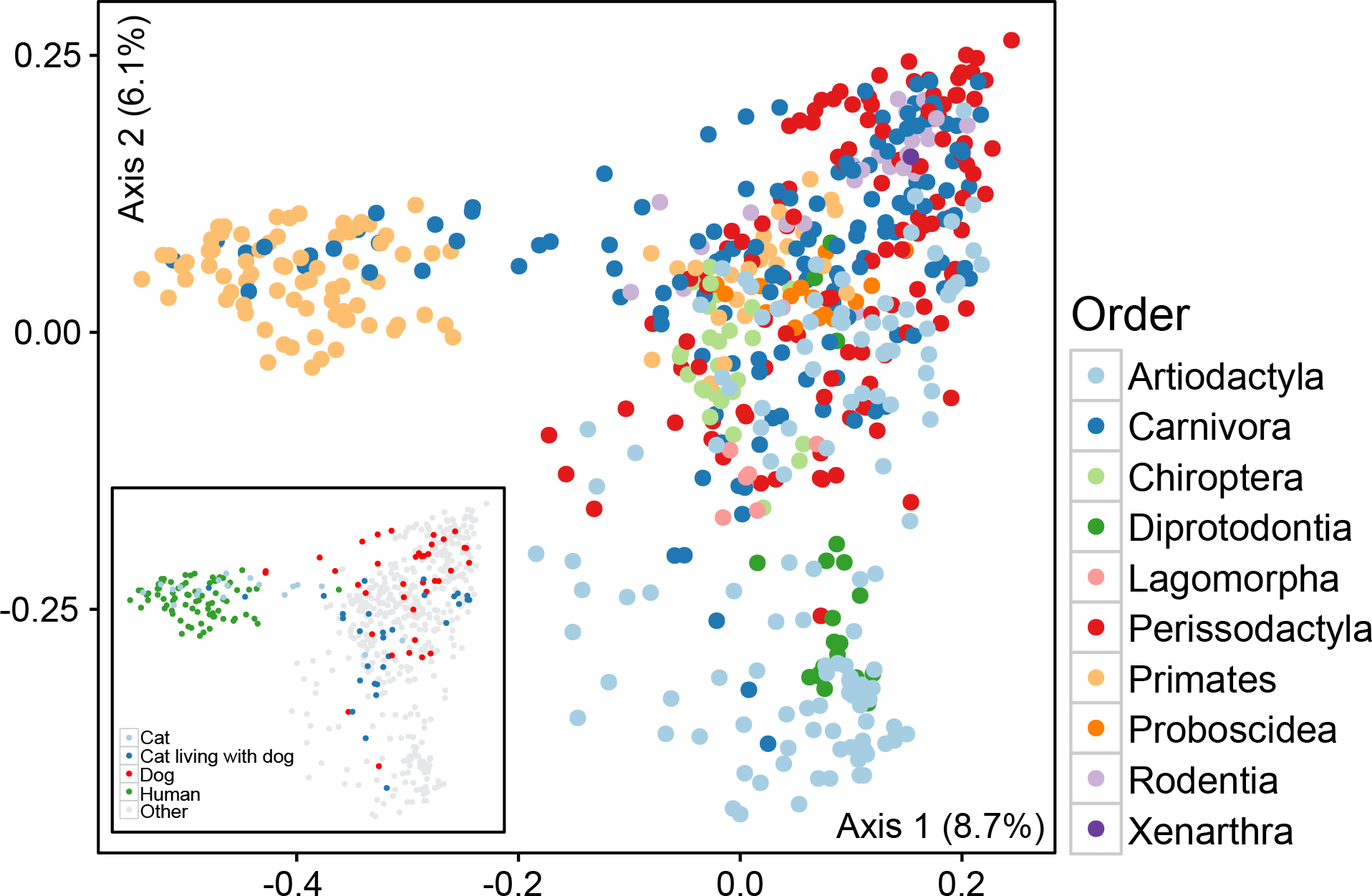
Ordinations (PCoA) generated by using the Bray-Curtis dissimilarity metric for each of the three body locations sampled. Samples are colored according to mammalian order. Inset: Ordination coloured according to human and non-human samples.

**Figure 2:**
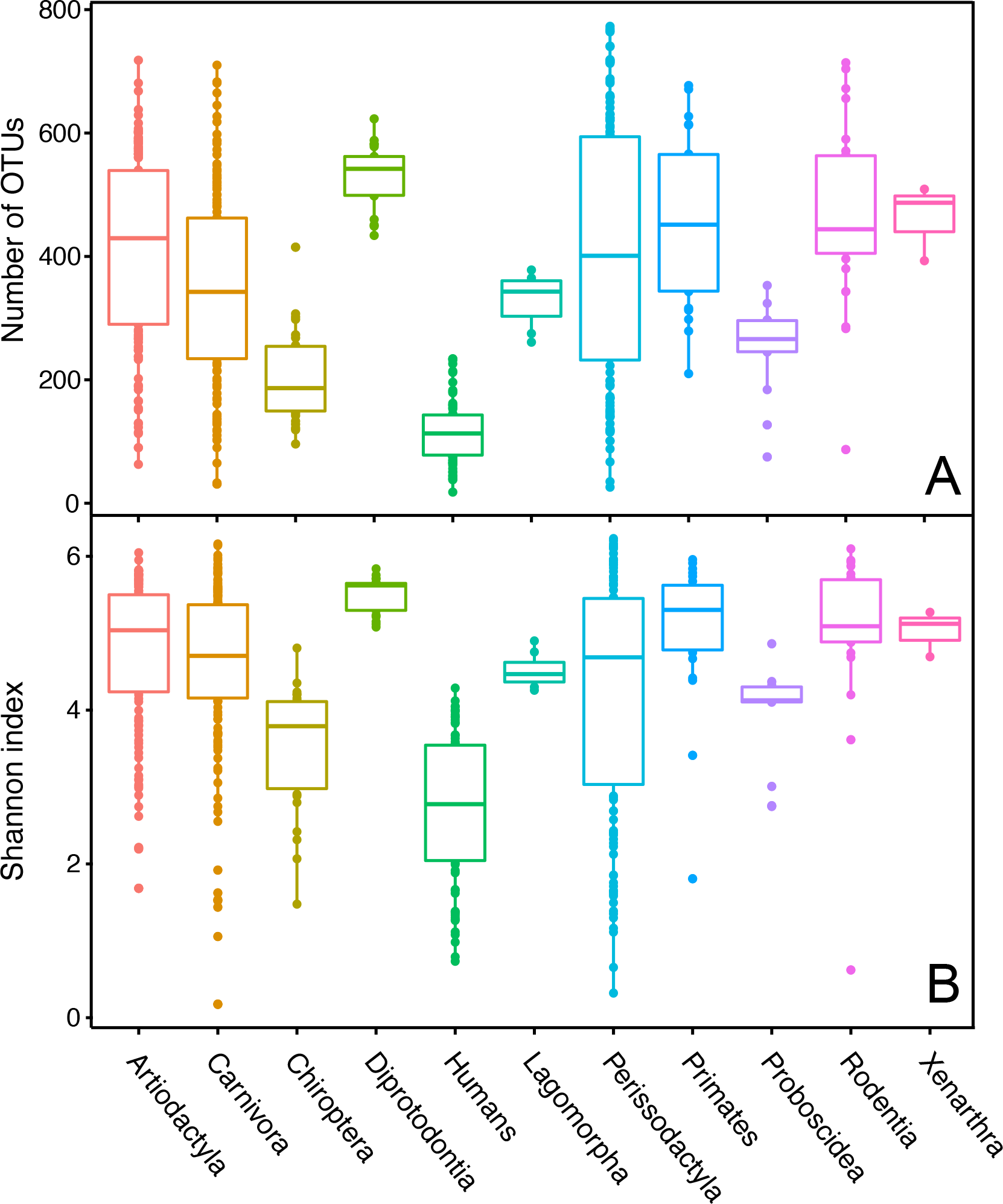
Boxplot of Shannon indices for 10 mammalian orders and humans.

**Figure 3:**
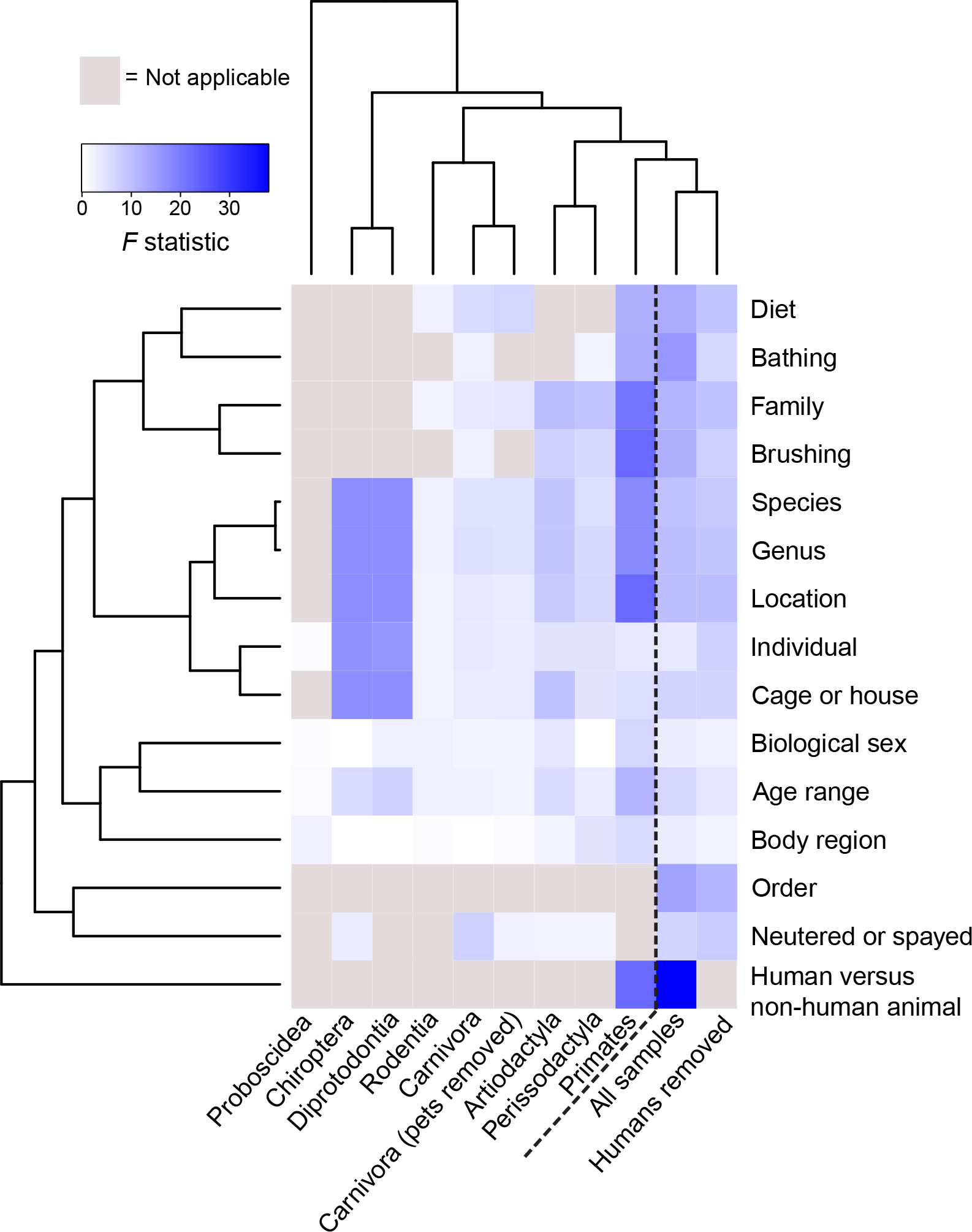
Heatmap summarizing the significant metadata factors correlating with the observed skin microbiota for sampled individuals from mammalian orders. Categories with higher PERMANOVA *F* statistics have higher variation in community dissimilarity. Grey regions of the heatmap represent categories that do not apply. Samples and categories are clustered according to Bray-Curtis distances.

By analyzing all samples together, random forest modelling identified that human and animal samples could be distinguished correctly 98.5±1.2% of the time. The OTUs that contributed most to the model include *Corynebacterium* (2.0%), *P. acnes* (1.2%), *Moraxellaceae* (1.2%), and *Macrococcus* (0.8%). These organisms were all within the top 10 most abundant OTUs in a dataset of all samples. A single female human back was grouped with the majority of the animal samples because of elevated abundances of *Luteimonas*, *Planomicrobium*, and *Planococcaceae*. The animals that were incorrectly classified were house pets, which had elevated levels of *Corynebacterium* and *P. acnes*. The specific house pets that grouped predominately with humans lived exclusively indoors. When all pets were removed from the dataset, humans could be distinguished from animals 99.8% of the time, which is 78.2-fold better than expected by chance.

Studies to optimize skin sampling methodology should be conducted to determine if there is a more optimal protocol to accurately sample the mammalian skin microbiota. Mammals were associated with OTUs that are traditionally associated with environmental environments, for example soil (Supplementary Figure 2), indicating that a washing or hair clipping step to remove external transient organisms may result in more accurate sample collection of the “true” skin microbial community. To date, the protocol for terrestrial mammals has generally not involved a washing step to remove external debris, although it was used previously in an axilla culturing study that was conducted in the 1950s (5). Amphibian skin microbiota studies have demonstrated that the microbial community in rinse water differed significantly from the community on the rinsed skin (35), and is widely adopted in amphibian research (35, 40–46). Although the current study observed that the external surface of a human differs significantly from all other mammals, more methodological testing needs to be done to determine if a wash step or haircoat clipping would provide a more accurate representation of the mammalian skin microbiota.

### Taxonomic order and host geographic location influence the mammalian skin microbiota

The effects of mammalian taxonomy, body region, and location were analyzed to elucidate whether these factors contributed to the detected skin microbiota. Mammalian order had the strongest association with the observed variation among animal skin communities (PERMANOVA; F9,502 = 11.3, p < 0.001; Figure 3), which was followed by geographic location (PERMANOVA; F4,507 = 9.6, p < 0.001; Figure 3). Random forest modelling was also conducted on a dataset of only animals to determine how well intrinsic factors (e.g., host taxonomy) and extrinsic factors (e.g., location) could be classified. Animals could be classified best according to their taxonomic order. This model was correct at classifying animals into their corresponding order 87.8±5.0% of the time, which is 5.9-fold better than expected by chance. Lower taxonomic orders, such as family (86.1±3.9%), genus (84.4±4.7%), and species (83.4±6.7%) were progressively classified less accurately. This weaker classification ability may be in part due to smaller number of samples per group for training the model.

The ability to classify accurately from specific locations may be in part due to the soil that is present in a given habitat. Indeed, a study that analyzed the similarity of skin bacterial communities between salamanders and their environment noted that certain taxa were shared between the skin microbiota and the abiotic environment (47), in part due to contact with forest litter. A previous study noted that the host was the most important factor influencing the skin microbiota of amphibians, whereas geographic location was the second most important factor (40), which aligns closely with both the PERMANOVA and random forest model findings from this study.

Other factors that have been demonstrated to influence the human skin microbiota, such as individuality, biological sex, and body region, exhibited less of an effect on animals. Both taxonomic order and geographic location were classified more accurately than biological sex (65.2±4.5%), body region (39.9±6.0%), or individual animal (36.7±36.7%). This inability to classify individual is in contrast to human studies that have shown that individuality is one of the most important factors influencing the human skin microbiota. However, many of these studies used >15 samples per individual (12, 48, 49). It is therefore still possible that animals’ skin microbiota are relatively unique among individuals, but this cannot be observed with only three samples per animal.

To address whether biological sex influenced the skin microbiota within a species, cat (n=48), dog (n=35), and horse (n=68) samples were analyzed because they contained a relatively large number of sampled individuals and a balanced biological sex split. Biological sex was not a significant factor for any of these species (PERMANOVA; Cat: *F*_1,15_ = 1.15, *p* = 0.20; Dog: *F*_1,11_ = 0.79, *p* = 0.77; Horse: *F*_1,20_ = 0.94, *p* = 0.44). All of the domestic cats and dogs were neutered or spayed, so this would not have an influence on the analysis. Although the horses had both neutered/spayed and intact individuals, whether the horse had been neutered or not did not exhibit a significant difference on the skin microbiota (PERMANOVA; *F*_1,20_ = 1.09, *p* = 0.34). The only animal where biological sex explained the most variation was the red kangaroo (PERMANOVA; *F*_1,16_ = 2.21, *p* = 0.002), which was also analyzed because this species exhibited a visual split between males and females in an ordination (PCoA;Supplementary Figure 3). Larger variations in the microbiota among different body regions were observed within Perissodactyla (PERMANOVA; F2,121 = 4.26, p < 0.001; Figure 3) and Proboscidea (PERMANOVA; F2,12 = 2.38, p = 0.02; Figure 3) than within other orders. The overall low effect from body region is likely due to the body regions sampled. The back, inner thigh, and torso are all covered with hair. A previous study on dogs demonstrated that fur-covered regions had higher species richness and diversity (25) compared to mucosal surfaces. Therefore, sampling mucosal surfaces would be expected to result in more distinct differences between body regions within a species.

To ensure that the importance of the host’s taxonomic order on skin microbiota was not overly influenced by orders with fewer samples and locations, the three orders with a large number of samples, Artiodactyla, Carnivora, and Perissodactyla, were analyzed with all other orders removed. Each of these orders had samples sourced from 6-8 different locations. Removing orders with fewer samples increased the influence of order (PERMANOVA; *F*_2,385_ = 15.1, *p* < 0.001) and decreased the impact of geographic location (PERMANOVA; *F*_3,384_ = 8.7, *p* < 0.001). Therefore, the effect on microbial communities exhibited by the mammalian host exists despite varying geographic locations, and cannot be fully attributed to certain species only being sampled in a single location.

### Pets and humans

Although the majority of animals possessed skin microbial communities that were distinct from humans, a subset of pets grouped with humans in ordination space (Figure 1). In particular, of the 17 pet samples that grouped with humans, 15 were from indoor housecats, whereas the remaining two samples belonged to the backs of dogs that were frequently bathed and groomed (Supplementary Table 2). None of these 17 pet samples belonged to animals that had been exposed to antibiotics in the past six months. In total, 75% of these samples belonged to animals that were owned by humans who were also sampled for this study. All cats with similar microbial communities to humans had at least two of the three sampled body locations possess this “human” community composition (Supplementary Table 2), whereas the two dogs only possessed the human microbial community on their backs. The remaining 11 dogs had similar communities to the other animals, as did all cats that lived outdoors on farms and a single cat that lived in the city, without a dog (Figure 1 inset). Interestingly, 11 of 12 indoor cats that lived with a dog possessed similar microbial communities to the other animals. It may be that owning a dog results in an influx of soil microbiota into the home, which in turn is transferred to the exclusively indoor cats through either contact with the built environment or personal contact. Indeed, several studies have shown that owning a dog shifts the human microbiota as well as built environment surfaces (50, 51). This study only sampled from 13 dogs and 19 cats (87 samples). Future studies might include a larger sample size of animals that would help further elucidate how owning a dog impacts other inhabitants within a household.

### Predicted functions of skin microbiota vary between human and animal samples

The predicted functions based on the prokaryotic clades were determined using FAPROTAX (52) and demonstrated that there were several conserved functions on mammalian skin (Figure 4). Many of these match functions predicted from human samples from the Human Microbiome Project consortium that underwent metagenomic sequencing (13, 53). Animal symbionts and human pathogens were expected because the samples were derived from mammalian hosts. Urea is a component of sweat and provides a nitrogen source, which could explain ureolysis as a predicted skin function, although not all animals sweat in hair covered regions (54). There were several functions that were significantly different between humans and animals. Humans had elevated levels of predicted manganese oxidation. Human sweat contains on average 100 ppb manganese, which would result in approximately 200-300 mg of manganese secreted each day (55). This concentration is low compared to other trace metal elements, such as zinc and copper (56). Manganese oxidation was predicted to occur from the core human OTU *P. acnes* (57). In contrast, animals had higher levels of predicted functions involved in the nitrogen cycle and single-carbon compound degradation. Methanol oxidation was attributed to the core OTUs affiliated with *Arthrobacter,* and methylotrophy with *Methylobacterium* OTUs, according to the FAPROTAX database. Nitrogen respiration was associated with numerous organisms, such as *Paracoccus* and *Pseudomonas*. In accordance with lower diversity (Figure 2), humans had a significantly lower number of predicted functions (34.2±8.4 compared to 51.8±9.4 in animals; *p* < 0.001). Animals may also have a higher number of predicted functions because they had more soil bacteria, which may have more annotated predicted functions than skin bacteria. Predicting functions based on taxonomy is the first step to elucidating how biochemical processes from skin microbiota are influencing host skin health. Future studies using metagenomic sequencing may help confirm which of these predicted conserved microbial functions are core to mammalian skin, versus functions that are variable amongst different host species.

**Figure 4:**
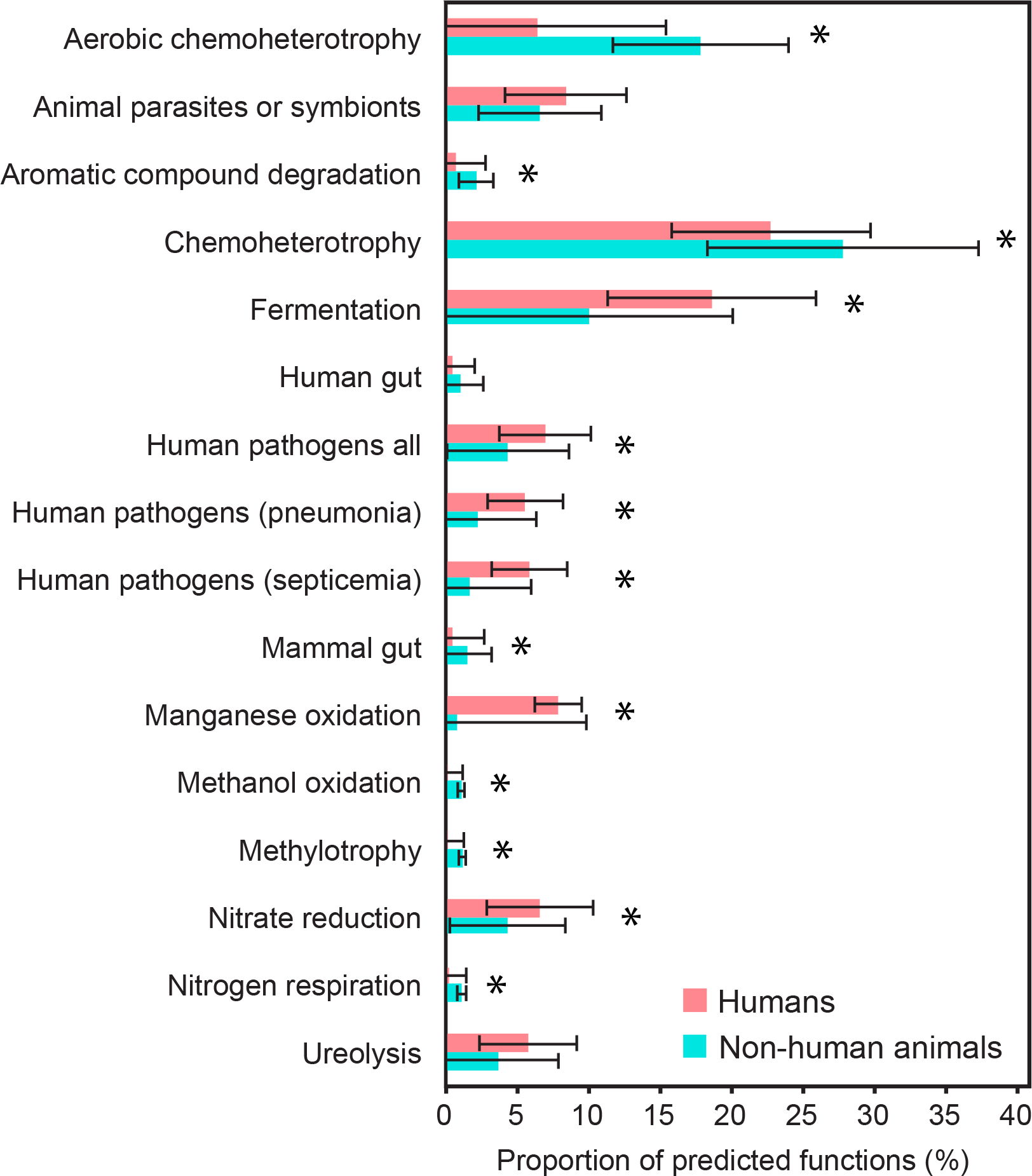
Barplot of predicted functions based on FAPROTAX database. Stars indicate*p* < 0.05 among mammalian and human samples after Bonferroni correction was applied. Error bars denote the standard deviation of animal (n=512) and human (n=77) samples.

### Phylosymbiosis is evident in the orders Perissodactyla and Artiodactyla

Phylosymbiosis postulates that closely related clades of animals will have more closely related microbial communities (58). This can be measured using normalized Robinson-Foulds values, which compares the congruence between two phylogenetic trees. A score of zero indicates the trees are identical, whereas a score of one indicates there is no congruence between the two trees. Previous studies have shown that shifts in microbial communities have matched host evolution within insects (58, 59), which were more apparent at the 99% OTU clustering threshold. In this study, comparisons were made at both the 97% and 99% prokaryotic OTU threshold, using Bray-Curtis, unweighted, and weighted UniFrac distances.

Comparing the known host mammalian phylogeny to dendrograms of the microbial communities for each host species supports the hypothesis that skin communities on animals from the orders Perissodactyla and Artiodactyla experienced shifts that match mammalian evolution. Perissodactyla exhibited phylosymbiosis with all thresholds and distance measures, because the only discrepancy in each test case was the microbial community of horses and Przewalski’s horses (Figure 5C). The split between the equestrian and rhinoceros clades cannot be attributed to differences in location, such as farm or zoo habitats, because the Przewalski’s horses were sourced from the Toronto Zoo. Although Artiodactyla (Figure 5A) possessed a relatively poor normalized Robinson-Foulds score of 0.71 (Table 3), it still demonstrated significant congruence with the host phylogeny and microbial dendrogram with both the Bray-Curtis and weighted UniFrac metrics. The largest discrepancy was noted with the sequences from the goats, which grouped with the giraffe and reindeer rather than the sheep. In addition, the host species did not group according to the geographic locations from where they were sourced. In contrast, the order Carnivora (Figure 5B) did not exhibit significant phylosymbiosis and the cat and dog clades did not have distinct microbial communities. This observation did not change when all cat and dog samples were removed from the dataset. Therefore, the microbial dendrograms were not being unduly influenced by household animals that undergo frequent grooming and spend the majority of time indoors. It is possible that phylosymbiosis may be more strongly observed within clades of animals that share similar diets or have similar management. All of the sampled animals within Perissodactyla and Artiodactyla were herbivores that graze on local grasses or hay. In contrast, the animals within order *Carnivora* had a more diverse diet between each species, such as herbivorous giant pandas, carnivorous lions, and pets that were fed an omnivorous diet. Similar to how diet influences the gut microbiota (60), it may be that the skin microbiota is impacted by diet, as has been shown for skin microbial communities of amphibians (41).

**Figure 5:**
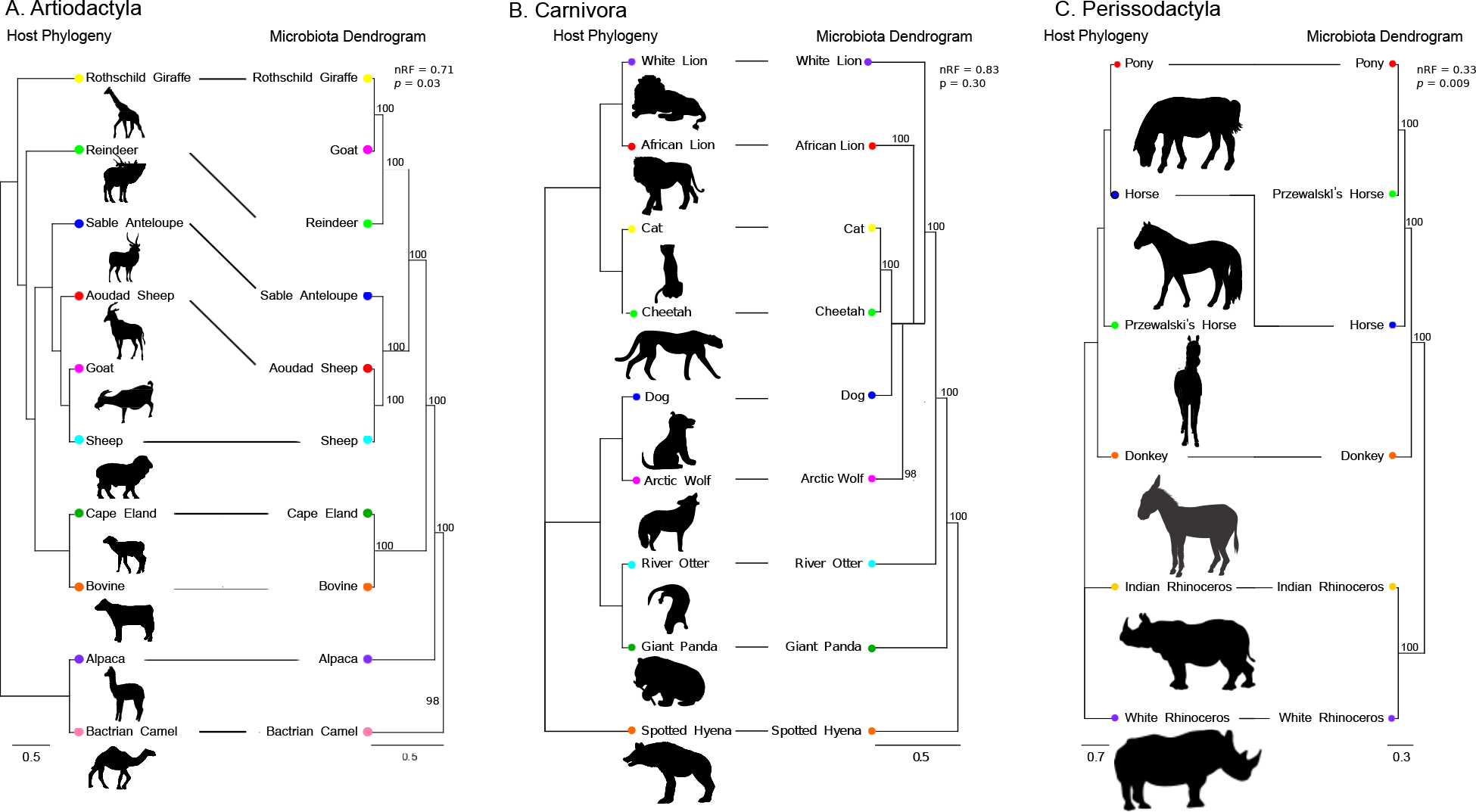
Microbiota dendrograms created using the Bray-Curtis distance metric and a 99% OTU threshold compared to the known host phylogenies of A: Artiodactyla, B: Carnivora, C: Perissodactyla. Congruences were measured using normalized Robinson-Foulds scores (nRF). Horizontal lines denote species that have concordance between the host phylogeny and microbial dendrogram. **Bootstrap values are located at the top-right of each split.** All images of are courtesy of Tracey Saxby, Integration and Application Network, University of Maryland Centre for Environmental Science, except for alpaca (Meaghan Mechler), bovine (John C. Fisher), goat (Jane Hawkey), and sheep (Tim Carruthers).

**Table 3:** Phylosymbiosis analysis of main mammalian clades. The normalized Robinson-Foulds scores were calculated at the 97% and 99% threshold. (BC: Bray-Curtis distance metric; nRF: normalized Robinson-Foulds score; UU: unweighted UniFrac distance metric; WU: weighted UniFrac distance metric). Significant normalized Robison-Foulds scores are starred.

| Clade | Distance metric | 97% threshold | 99% threshold |
| --- | --- | --- | --- |
| All samples | BC | 0.97 | 0.97 |
|  | UU | 1.00 | 0.93* |
|  | WU | 0.97 | 0.97 |
| All mammals:<br>humans removed | BC | 0.93* | 0.94* |
|  | UU | 0.96* | 0.94* |
|  | WU | 0.93* | 0.93* |
| Artiodactyla | BC | 0.71* | 0.71* |
|  | UU | 0.86 | 0.86 |
|  | WU | 0.71* | 0.71* |
| Carnivora | BC | 0.83 | 0.83 |
|  | UU | 1.00 | 1.00 |
|  | WU | 0.83 | 0.83 |
| Carnivora:<br>pets removed | BC | 1.00 | 0.75 |
|  | UU | 1.00 | 1.00 |
|  | WU | 0.75 | 0.75 |
| Perissodactyla | BC | 0.33* | 0.33* |
|  | UU | 0.33* | 0.33* |
|  | WU | 0.33* | 0.33* |

There was no significant phylosymbiosis observed when all mammalian orders and humans were analyzed, except for the unweighted UniFrac measure at the 99% threshold. Although in this single case animals could be matched significantly better than 100,000 randomized trees of the 38 species, there was very little congruence observed (Table 3), as indicated by the normalized Robinson-Foulds score of 0.93. When humans were removed from this dataset, congruence was increased modestly, although the host tree and bacterial dendrograms still exhibited little congruence. This can be explained by the significantly different microbial community that humans have, because human skin microbial community node was positioned near the root of the tree, instead of within the primate clade.

Although previous studies have been able to demonstrate phylosymbiosis, they did so under highly controlled laboratory conditions and with fecal samples (58). Skin represents a more transient environment that is influenced by shedding and contact with other surfaces. The animals in this study had several confounding factors, such as different locations and age (Supplementary Table 3). It is possible that if mammals were sampled at similar timepoints in their life history, and inhabited the same geographic location that a more distinctive congruence would be observed. Additionally, the potentially transient soil microorganisms that were abundant on mammalian skin may mask phylosymbiosis when all sequenced OTUs are being considered as a community in the phylosymbiosis analysis (Supplementary Figure 1). Future studies should potentially sample before and after washing the skin to observe how this treatment would influence the analysis. We postulate that reducing the number of transient auxiliary organisms from the environment would strengthen the finding of phylosymbiosis because the transient organisms that would not co-evolve with a host would be removed from the analysis. Phylosymbiosis between the skin microbial community and host would not be unexpected. Multiple studies have observed this phenomenon in the gut of mammals (60, 61). These findings illustrate that despite multiple confounding factors that would potentially mask phylosymbiosis, that it is still significantly observed in multiple mammalian clades. Further studies should determine if this finding is strengthened when the hosts within a clade experience equivalent extrinsic factors.

### Archaea are present on mammalian skin at low abundance levels

Archaea comprised only 6,509 of the total 6,550,625 non-rarefied sequences (0.001%; Supplementary Table 1). Several archaeal clades were present, such as the salt-tolerant *Halobacteria*, the methanogen *Methanobrevibacter*, and the ammonia-oxidizing *Thaumarchaeota* (Figure 6). Methanogens likely represent fecal contamination because *Methanobrevibacter* spp. are the dominant archaea present in the gut (62). However, *Halobacteria* and thaumarchaeotes, such as *Nitrososphaera*, have the potential to be resident skin microbiota. The *Halobacteria* are able to tolerate the salt concentrations from sweat (63), whereas putative ammonia-oxidizing archaea have been observed on human skin (64, 65).

**Figure 6:**
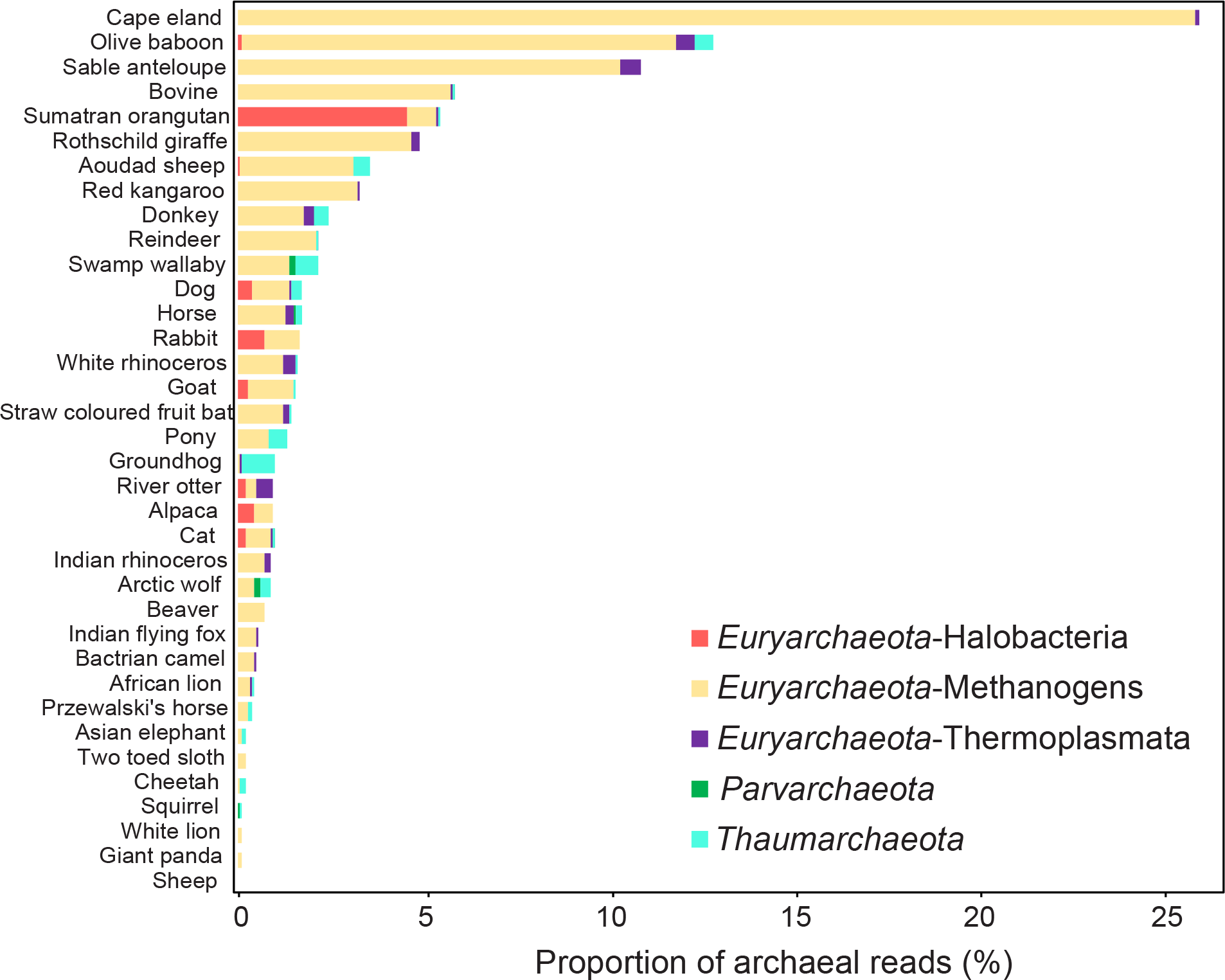
Barchart of archaeal sequence reads. The proportion of reads represents the total proportion of the 6509 archaeal reads. Each mammalian species was corrected by the number of samples collected to account for an unequal sampling depth.

Archaeal reads were disproportionately present on cape elands (26.1% of all archaeal sequences), olive baboons (12.9%), the sable antelope (10.9%), and bovine (5.8%). The methanogens from the phylum Euryarchaeota, were the dominant archaeal clade. This finding is expected because the animals with the most archaea were predominately ruminants. However, Thaumarchaeotes were significantly more predominant (*p* < 0.001) in groundhogs (0.9%), the swamp wallaby (0.6%), olive baboons (0.5%), and the pony (0.5%), providing further evidence to previous research that these organisms are a part of the skin microbiota at low relative abundance (64, 65).

The low relative abundance of archaea compared to bacteria in this study is likely an underrepresentation of the actual abundance because of primer mismatches to archaeal 16S rRNA genes (66). Indeed, the performance of the Pro341F/Pro805R primer was analyzed using TestPrime v. 1.0 on the SILVA database (Supplementary Table 4). Only 64.8% of archaeal 16S rRNA genes had zero mismatches to the primer used in this study. Disconcertingly, the ammonia-oxidizing thaumarcheotes only had 11.9% of taxa with zero mismatches, in contrast to 85.7% of all bacteria. When the number of mismatches was increased to two, 94.9% of all archaea, and 95.5% of thaumarcheotes were matched. A recent study on the gut microbiota of great apes that used both universal prokaryotic and archaeal specific primers determined that the distribution, diversity, and prevalence of archaea in mammalian gut samples is underestimated by up to 90% (66). Currently archaea-specific primers, namely Arch516F/Arch915R, offer more accurate representation of this domain. Therefore, this study provides evidence that archaea are present in relatively high abundance on cape elands, olive baboons, sable antelope, and bovine, which requires further examination with archaea-specific primers or metagenomics.

### Limitations

This study possesses several inherent limitations. The majority of the animals were collected based on opportunistic availability. For example, animals from the Toronto Zoo were sampled during routine veterinary checkups. The nature of sample collections resulted in an inability to collect an equal number of represents from each host taxonomic order and species. For example, the following species only had a single representative sampled: alpaca, beaver, pony, sheep, sable antelope, spotted hyena, swamp wallaby, and two-toed sloth. Although we recognize that no significant conclusions can be made about a single host animal within a species, these animals were included in the analysis to have the most in depth coverage of each mammalian order. This study represents an initial survey of the mammalian skin microbiota. Much work remains to be conducted within each species to determine intra-specific effects of individuality, body region, and biological sex.

The microbial communities on animals were sampled across an entire year and samples were frozen until DNA extraction. It is possible that the skin microbiota of outdoor animals may undergo seasonal shifts, especially between the relatively cold winter and warm humid summer in Canada; however, this cannot be tested using a single sampling time for each animal. Future investigations should sample the same individuals across a year to determine if changes in temperature and resulting skin secretion levels might exhibit an effect on the microbiota. Moreover, the significant difference in geographic location that was observed may be more pronounced if animals with greater geographic distance were sampled. All of the animals were sampled in Southern Ontario. Sampling the same species from multiple continents is hypothesized to result in more pronounced variations in communities according to location due to significant changes in extrinsic factors, such as soil microorganisms.

An inherent limitation of any amplicon study is biases that arise from primer selection. This study utilized the V3-V4 16S rRNA region. A larger target size has the potential for low level sequence error that can result in high OTU counts and corresponding alpha diversity. The relative abundances of common skin organisms, such as *Propionibacterium* and *Staphylococcus* may differ in studies that select another portion of the gene, for example the V1-V3 (67).

Although these biases have been discussed in detail elsewhere (67–70), it has been shown that the V3-V4 region can accurately represent skin microbial communities (71) and is therefore expected to influenced the results only minimally.

The rodents collected in this study were sourced from the wild and deceased. Although these samples still grouped with the remaining live animals (Figure 1), high skin microbial diversity (Figure 2) may be in part related to initial changes in skin community from decomposition, which have been shown to progress in a clock-like manner (72). Deceased rodents were collected during the first day of death and did not have any visible injuries that would result in internal microorganisms from the gastrointestinal tract contaminating the skin.

## Conclusion

Human samples were dominated by *S. epidermidis*, *Corynebacterium*, and *P. acnes*. In contrast, other animals were significantly more diverse and have higher levels of soil OTUs that likely represent transient organisms from their enclosure or natural habitat. These findings demonstrate that human skin is distinct, not only distinct from other primates, but from all ten mammalian orders sampled. Given the recent evolutionary divergence of humans as distinct species from other non-human primates, these results suggest that modern human practices, such as living within a built environment, wearing clothing, and washing with soap, have strongly impacted the diversity and composition of the skin microbiota that can be sampled with sterile swabs.

## Materials and Methods

### Ethics

The study was approved by the Office of Research Ethics at the University of Waterloo (A-15-06). The following minimally invasive procedures were conducted in accordance with the approved documentation and no animals were harmed throughout this study.

### Sample collection

Species from ten orders of the class Mammalia were sampled to characterize the distribution of microorganisms on skin (Table 1). Both males and females were included for each species, when available, to account for variations in hormone levels and secretions that are known to affect microbial communities (33). A spectrum of habitats and hygiene practices were also included, ranging from frequent grooming of pets and farm animals, to animals in captivity and the wild. Complete information on the biological sex, age, diet, location, health history, grooming, and exposure to antibiotics were collected (Supplementary Table 3). The inclusion of animals that had been exposed to antibiotics in the previous six months was minimized (i.e., 32 animals; 10 in previous 2 months), and did not influence diversity within a species.

Animals were sampled from multiple locations in Southern Ontario from November 2015 to September 2016: The African Lion Safari, Kitchener-Waterloo Humane Society (KWHS), Toronto Zoo, pet owners, and from local farms sourced from the University of Guelph. Animals from the Toronto Zoo and African Lion Safari were sampled when they were brought in for regular husbandry practices. Additional companion animals were obtained from volunteers who were recruited by word of mouth. The KWHS supplied wild animals that were collected by KWHS staff within 24 hours of death; the specimens were stored in plastic bags in a −20°C freezer until sampled.

The back, torso, and inner thigh regions of 177 non-human mammals were collected using sterile foam swabs (Puritan) according to a previously published protocol (32). In addition, we included data from 77 equivalent samples (the right and left inner thigh were sampled from each participant) from 20 human participants from a previous study that were sampled from November 2015-February 2016 (32) in the analysis for comparison purposes, for a total of 589 samples. These regions were chosen to capture both moist and dry regions and avoid sensitive areas that may cause distress. Skin was swabbed by moving aside hair or fur with gloved hands to expose the skin. While applying moderate pressure, the skin was swabbed 10 times in a forwards and backwards motion. The swab was rotated and repeated in adjacent areas for a total of 40 strokes per swab. When the area was complete, the samples were returned to their initial plastic storage container and frozen at −20 °C until further use. All volunteers and veterinary technicians were trained with a detailed protocol to ensure sample collection consistency.

### Sample preparation

All DNA extractions, PCR protocols, and Illumina sequencing were conducted according to a previously published protocol (32) to enable comparisons between the human and nonhuman samples. In brief, DNA was extracted using the PowerSoil-htp 96 Well DNA Isolation Kit (MO BIO Laboratories) and stored at −20°C until further use. The V3-V4 region of the 16S rRNA gene was amplified using the Pro 341Fi (5’-CCTACGGGNBGCASCAG-3’) and Pro 805Ri (5’-GACTACNVGGGTATCTAATCC-3’) primers (73). Each amplification was performed in triplicate to minimize potential PCR bias from low biomass samples (74), and was conducted in a PCR hood that was UV treated for 30 min after having undergone a treatment with UltraClean Lab Cleaner (MoBio) to remove DNA, RNA, DNase, and RNAses (75).

Six “run control” samples consisting of human, zoo, pet, and wild animal samples were included in each of the three runs, confirming the absence of detectable run bias (Supplementary Figure 4). The low diversity observed in human samples (Figure 2) was not due to variations in Illumina run sequencing because the 37 non-human animal samples included in the first lane possessed the same diversity levels as samples from the same species that were sequenced in other lanes (data not shown). The no-template, DNA extraction kit, and sterile swab controls were analyzed for contaminants after sequence processing (Supplementary Table 5), and were determined to not influence the study (refer to Supplementary Information for in-detail analysis of controls). To reduce the known impact from well to well cross-contamination (75), all samples were randomized. This ensured that samples from the same animal were distributed across multiple plates and MiSeq runs, as were samples from within a mammalian species or order. Observed influences, including host taxonomy and geography, cannot be due to these groups of samples being situated proximally within the same extraction or PCR plate.

### Processing of sequence data

Raw DNA sequence reads were processed using the same open source bioinformatics pipeline described previously (32) that was managed by Automation, eXtension, and Integration Of Microbial Ecology: v. 1.5 (76). PANDAseq v. 2.8 (77) generated paired-end sequences using the default parameters of a 0.9 quality threshold, a minimum sequence overlap of 10 bases, and a minimum read length of 300 bases. Quantitative Insights Into Microbial Ecology (QIIME) v. 1. 9.0 (78) was used to analyze sequence data, which underwent *de novo* clustering and chimera/singleton removal at both 97% and 99% cluster identity using UPARSE (79). PyNAST v. 1.2.2 (80) was used to align 16S rRNA gene sequences. Subsequently, RDP v. 8.1 (81) assigned prokaryotic taxonomy using Greengenes database v. 13.8 (82). Samples were rarefied to 1654 sequences in the dataset that contained all samples (Supplementary Table 1). Analyses such as alpha diversity and PERMANOVA underwent 1000 iterations of rarefication to avoid underrepresenting diversity. Rarefication plots demonstrated that conducting multiple rarefications to determine diversity prevented a loss in diversity levels (data not shown). Other mammalian skin microbiome studies have used a similar level of sequences to analyze their data (25, 27, 83, 84).

### Negative control analysis

The no-template, DNA extraction kit, and sterile swab controls were analyzed for contaminants after sequence processing (**Supplementary Table 5**). A total of 3 of 4 kits controls, 4 out of 5 sterile swabs, and 67 out of 69 no template PCR controls contained fewer reads than all other samples. The sterile swab and kit control that contained a higher number of sequences were processed with different kits, implying that contamination from an adjacent well may have impacted this kit control (59), instead of an inherent contamination within the DNA extraction reagents (the contaminated kit control was processed in a plate with a clean sterile swab, and vice versa). The most abundant kit control contaminant was related to the *Neisseriaceae*, at 48.7% abundance in the control sample. This OTU was present in ~27% of samples in this run, the majority of which were cats. Indeed, cat #136 had a very high number of *Neisseriaceae* sequences (~42,000), and was located adjacent to the kit control well. It is therefore hypothesized that this particular kit control’s high contamination was from adjacent well via cross-contamination instead of from a source that would impact all samples, such as kit reagents, implying that there was no significant impact on all samples. Additionally, one of the contaminated no template PCR controls was dominated by an OTU affiliated with *Rhodocytophaga* (36.2%), which had only a single read in one animal sample included in the study. To reduce the known impact from well to well cross-contamination (75), all samples were randomized. This ensured that samples from the same animal were distributed across multiple plates and MiSeq runs, as were samples from within a mammalian species or order. Observed influences, including host taxonomy and geography, cannot be due to these groups of samples being situated proximally within the same extraction or PCR plate.

The following 19 animal swabs were removed in the mammal dataset due to failure to amplify: eight cats, two beavers, and one each of river otter, cape eland, white rhinoceros, cheetah, horse, dog, Indian flying fox, and reindeer samples. These unamplified samples represent 3.6% of total mammalian samples. There was a disproportionate number of cat samples requiring removal, which may be due to several factors, such as pet owners sampling more lightly on cats resulting in insufficient sample collection. If the swabs were not pressed firmly against the animal’s skin, it is possible that only a small number of microorganisms were collected that were below the sequencing detection limit. Alternatively, cats may possess lower overall skin microbial abundances.

### Statistical analyses

The majority of statistical analyses were conducted using the same programs and software version numbers as described previously (32). In brief, alpha-diversity of all samples was measured with the following QIIME commands: multiple_rarefaction.py, alpha_diversity.py, collate_alpha.py, and compare_alpha_diversity.py. Subsequently, the 42 metadata categories were compared using the Bonferroni correction to avoid false positives due to testing a high number of hypotheses.

Beta diversity was visualized using ordinations generated with the Bray-Curtis distance metric. These figures were created in RStudio (85) with the phyloseq (v. 1.14.0) and ggplot2 (v. 2.1.0) packages. Beta diversity was measured using permutational multivariate analysis of variance (PERMANOVA) with the adonis function from the vegan package (v. 2.4-0) in R. Using 1000 permutations, the percent variation explained by each metadata category was calculated and visualized in a heatmap using ggplot2, vegan, Heatplus 2.16.0 (86), and RcolorBrewer v. 1.1.2 (87).

The functions from the prokaryotic clades were predicted using Functional Annotation of Prokaryotic Taxa v.1.0 (FAPROTAX) (52). This conservative algorithm currently matches 80 functions, such as fermentation and methanogenesis, against 7600 functional annotations of 4600 prokaryotic taxa.

### Phylosymbiosis analysis

The phylosymbiosis analysis of skin microbiota profiles and host phylogeny was adapted from a previously described protocol (58). Downloaded COX1 sequences were aligned with Muscle v. 3.8.31(88) and edited by removing gap positions and 5’/3’ end overhangs with Jalview v. 2.9 (89). The final edited alignment was created using RaxML online Blackbox server v. 8.2 (90). All mammalian host trees were verified to be in concordance with well-established mammalian phylogenies (91–95).

Microbiota dendrograms were constructed using the QIIME v. 1.9.0 jackknifed_beta_diversity.py command. Species were rarefied to the highest possible sequence count that included all species within the specific taxonomic ranking. This resulted in a rarefication of 1900 sequences for all mammals, 9,100 for Artiodactyla, 25,7000 for Carnivora, and 37,500 for Perissodactyla. Rarefication was conducted 1000 times and a consensus tree built to correct for the low number of rarefied sequences. Each of the above mammalian clades had bacterial consensus dendrograms created at 97% and 99% OTU identity threshold using Bray-Curtis, weighted and unweighted UniFrac distance metrics.

Congruencies between host phylogenies and microbial dendrograms were determined using the ape R package (96). Normalized Robinson-Foulds scores were calculated to quantify congruence (97). The significance of this score was determined by constructing 100,000 randomized trees with identical leaf nodes to the bacterial dendrograms and comparing each to the host phylogeny to calculate the number of stochastic dendrograms with equivalent or better Robinson-Foulds scores.

## Data Availability

The sequence data associated with all mammal samples are available in the Sequence Read Archive (SRA) under BioProject ID PRJNA385010. The sequence data for all human participants (published previously) are available in the Sequence Read Archive (SRA) under BioProject ID PRJNA345497.

## Acknowledgements

We thank Rahgavi Poopalarajah and Mayar Zawawi for sample kit preparation in addition to Elena Dybner for sample kit preparation and data entry. We also thank Katja Engel for assistance with Illumina library preparation. We thank Charles Gray and his team from the African Lion Safari, Dr. Diego Gomez from the University of Guelph, Andrea Dada, Dawn Mihailovic and the veterinarian technicians from the Toronto Zoo, and Amanda Hawkins and Joe Growden from the Kitchener-Waterloo Human Society. We would also like to thank Heather Cray, Emilie Spasov, and all farmers and pet owners who participated in the study.

## Supplementary Figure Legends

**Supplementary Figure 1:**
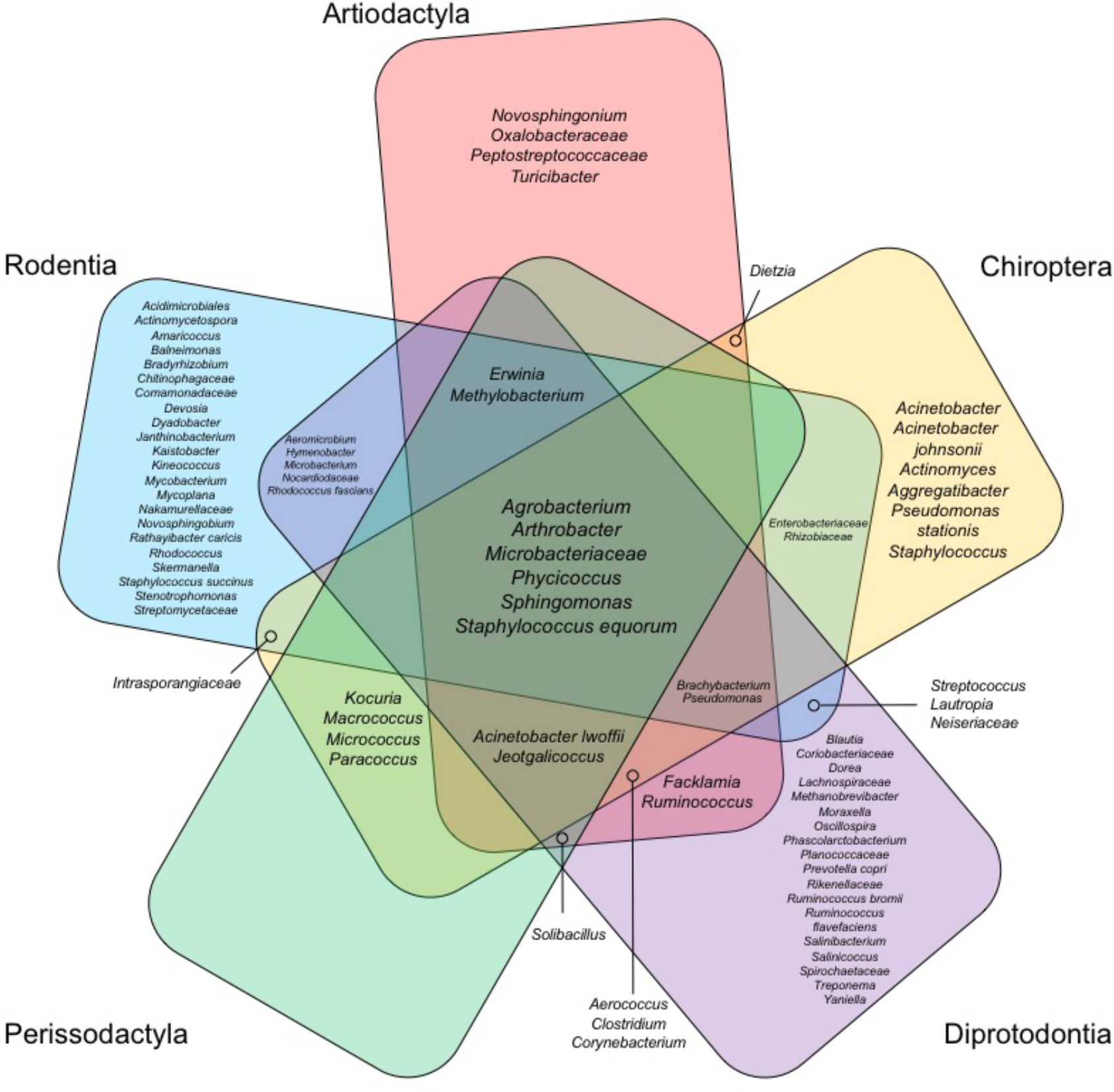
Venn diagram of core OTU analysis. A core OTU was defined as being present in >90% of samples in a designated category. The five mammalian orders were included that had multiple species, and did not have animals that typically inhabit indoors, such as humans, cats, and dogs. The most resolved taxonomic ranking for each OTU was included.

**Supplementary Figure 2:**
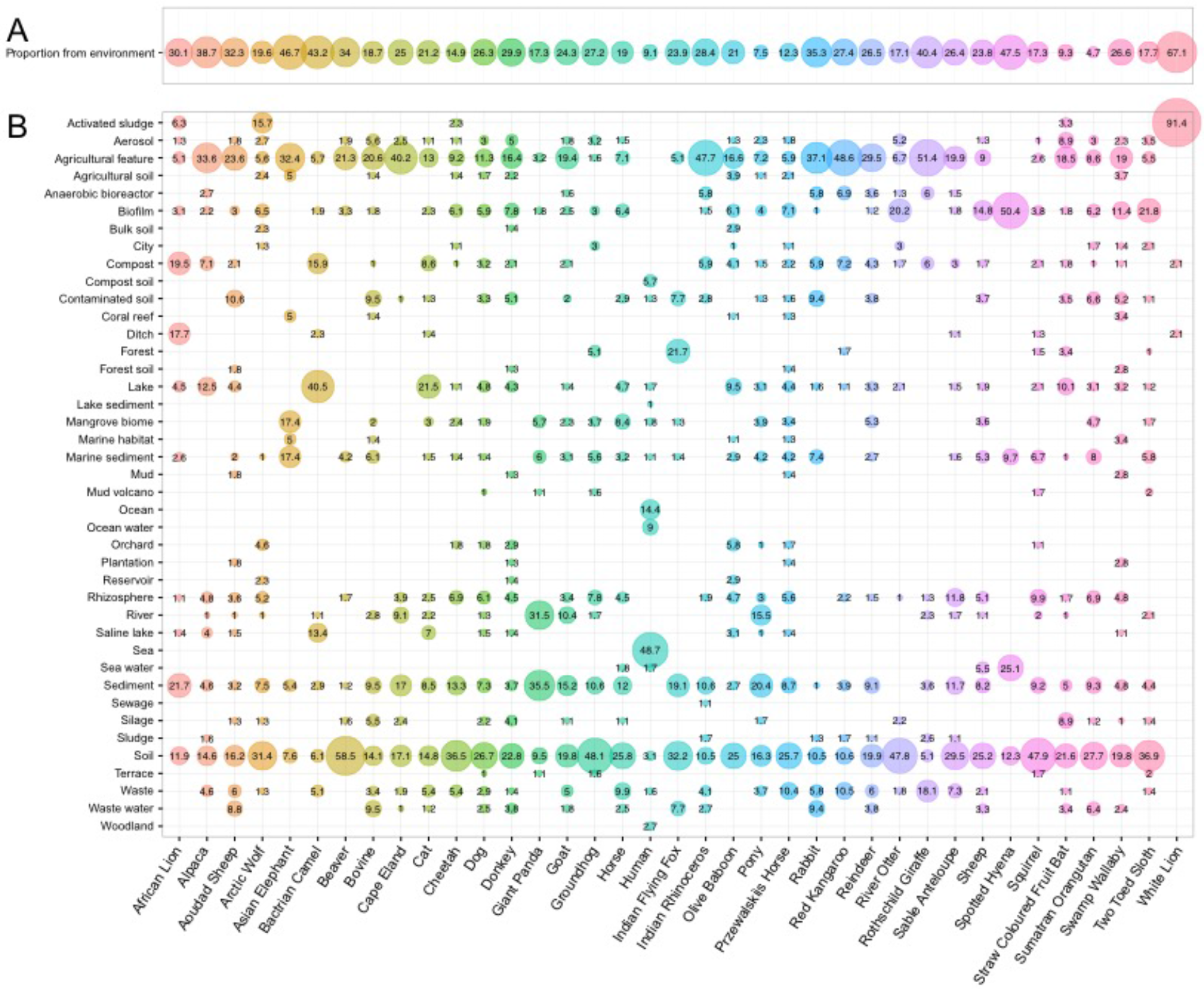
Bubbleplot of the proportion of OTUs associated with a non-skin environment for each mammalian species, according to a SeqEnv analysis. A. Proportion of total sequences that were not associated with skin. B. Distribution of non-skin associated sequences across environmental habitats. Only environments present >1% relative abundance are shown.

**Supplementary Figure 3:**
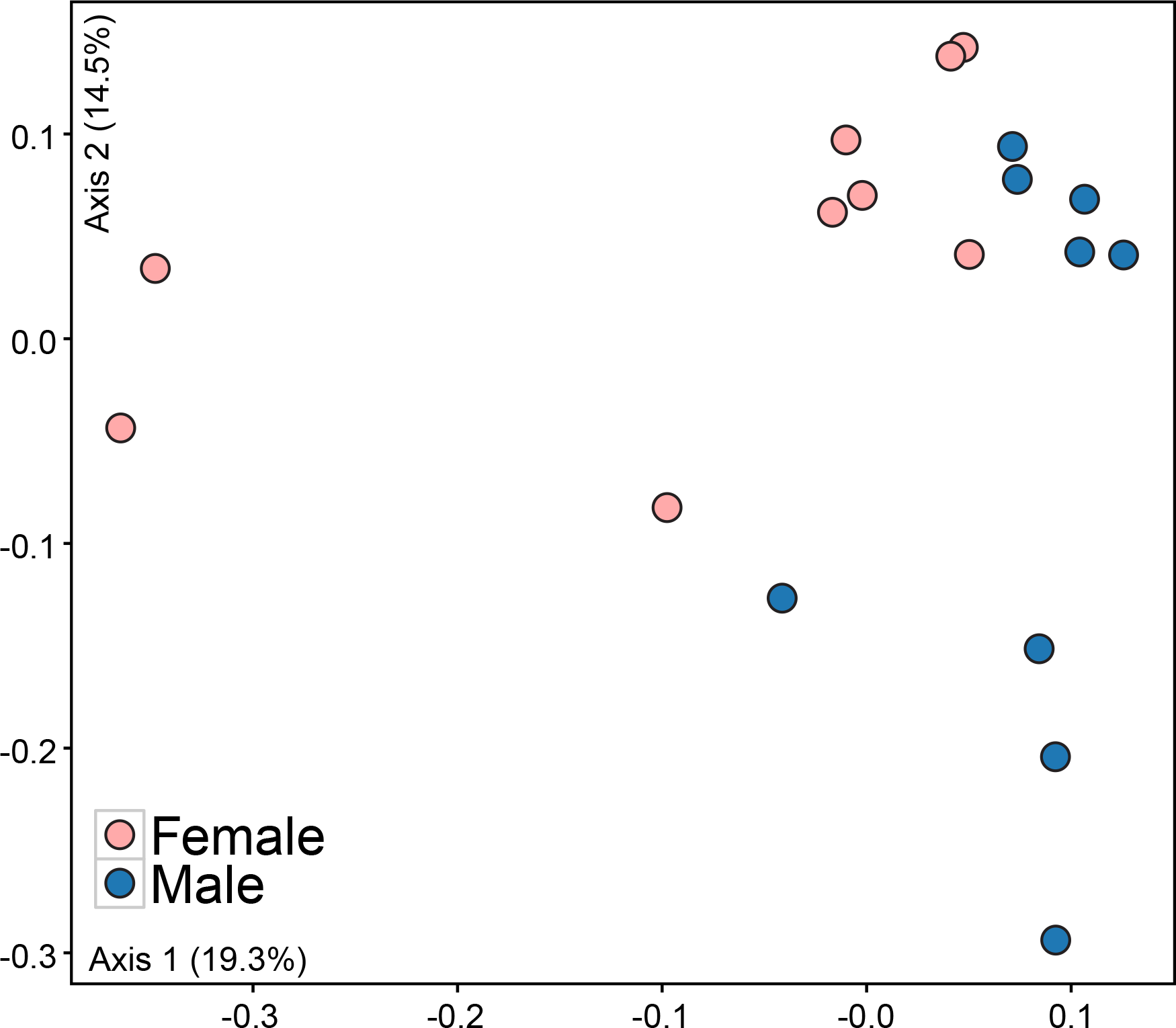
Ordination (PCoA) generated by using the Bray-Curtis dissimilarity metric for each of the three body locations of red kangaroos. Samples are colored according to biological sex.

**Supplementary Figure 4:**
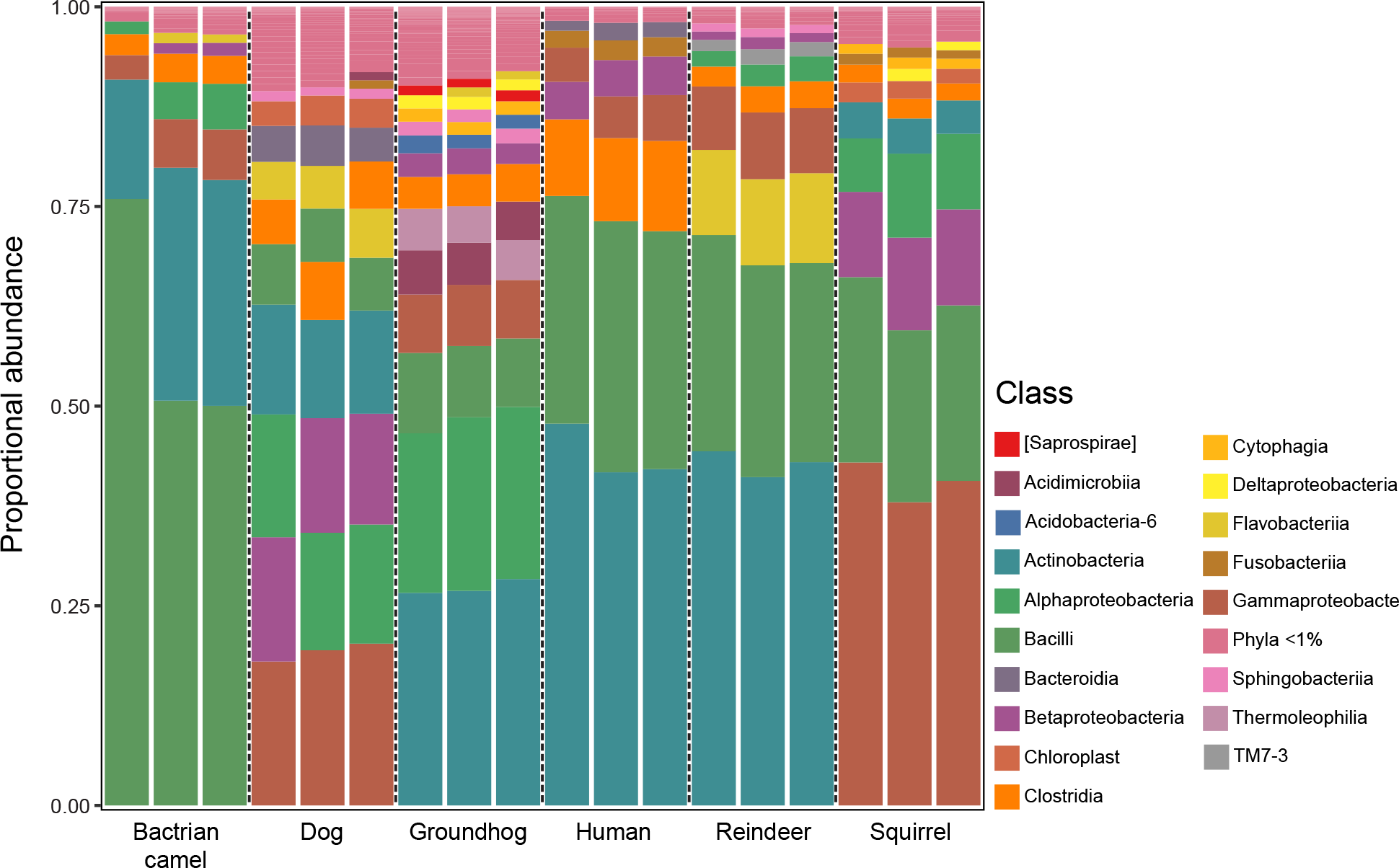
Taxaplot of six “run control” samples included in each of the three MiSeq runs. OTUs present >1% relative abundance were visualized.

## Supplementary Table Legends

**Supplementary Table 1:** OTU table organized in multiple tabs. Tab 1 contains the original rarefied OTU table, tab 2 contains the rarefied table with only mammalian samples, and tab 3 is a condensed table containing only archaeal reads. Sample names are explained in Table S3.

Dataset attached separately.

**Supplementary Table 2:** Table containing the animal samples that had similar microbial communities to humans.

| Animal ID # | Species | Owner is also in the study? | Number of samples grouped with humans | Body regions that grouped with human |
| --- | --- | --- | --- | --- |
| 52 | Cat | Yes | 3 | Back, inner thigh, torso |
| 54 | Cat | Yes | 2 | Back, torso |
| 55 | Cat | Yes | 2 | Back, inner thigh |
| 56 | Cat | Yes | 2 | Inner thigh, torso |
| 57 | Dog | Yes | 1 | Back |
| 58 | Cat | Yes | 3 | Back, inner thigh, torso |
| 69 | Dog | No | 1 | Back |
| 70 | Cat | No | 3 | Back, inner thigh, torso |

**Supplementary Table 3:** Metadata table containing all survey responses, PCR setup information, and sample codes. Animal samples names are coded as follows: A = Animal, # = the animal subject number, B = Back, I = Inner thigh, T = Torso. AL= African Lion, AP=Alpaca, Aoudad Sheep=AS, Arctic Wolf = AW, Asian Elephant = AE, Bactrian Camel = BC, Beaver = BE, Bovine = BV, Cape Eland = CE, Cat = C, Cheetah = CH, Dog = D, Donkey = DK, Giant Panda = GP, Goat = G, Groundhog = GH, Horse = H, Indian Flying Fox = IFF, Indian Rhinoceros = IR, Olive Baboon = OB, Pony = P, Przewalski’s Horse = PH, Rabbit = RB, Red Kangaroo = RK, Reindeer = R, River Otter = RO, Rothschild Giraffe = RG, Sable Antelope = SA, Sheep = S, Spotted Hyena = SH, Squirrel = SQ, Straw Coloured Fruit Bat = FB, Sumatran Orangutan = SO, Swamp Wallaby = SW, Two-Toed Sloth = TS, White Lion = WL, White Rhinoceros = WR. Human sample code names are coded as follows: H01-H10 signify each couple, whereas A-B differentiates individuals within a couple, 09 = torso, 10 = back, 12 = left inner thigh, 13 = right inner thigh.

Dataset attached separately.

**Supplementary Table 4:** TestPrime comparison of Pro341F/Pro805R primer mismatches to archaea, thaumarchaeotes, and bacteria.

| # mismatches | % archaea covered | % thaumarchaeotes covered | % bacteria covered |
| --- | --- | --- | --- |
| 0 | 64.8 | 11.9 | 85.7 |
| 1 | 89.0 | 93.2 | 94.6 |
| 2 | 94.9 | 95.5 | 96.1 |

**Supplementary Table 5:** Non-rarefied OTU table of the 77 negative controls included in the study.

Dataset attached separately.

